# Computational investigation of the effect of reduced dynein velocity and reduced cargo diffusivity on slow axonal transport

**DOI:** 10.1101/2022.10.14.512302

**Authors:** Ivan A. Kuznetsov, Andrey V. Kuznetsov

## Abstract

Contributions of three components of slow axonal transport (SAT) were studied using a computational model: the anterograde motor (kinesin)-driven component, the retrograde motor (dynein)-driven component, and the diffusion-driven component. The contribution of these three components of SAT was investigated in three different segments of the axon: the proximal portion, the central portion, and the distal portion of the axon. MAP1B protein was used as a model system to study SAT because there are published experimental data reporting MAP1B distribution along the axon length and average velocity of MAP1B transport in the axon. This allows the optimization approach to be used to find values of model kinetic constants that give the best fit with published experimental data. The effects of decreasing the value of cargo diffusivity on the diffusion-driven component of SAT and decreasing the value of dynein velocity on the retrograde motor-driven component of SAT were investigated. We found that for the case when protein diffusivity and dynein velocity are very small, it is possible to obtain an analytical solution to model equations. We found that, in this case, the protein concentration in the axon is uniform. This shows that anterograde motor-driven transport alone cannot simulate a variation of cargo concentration in the axon. Most proteins are non-uniformly distributed in axons. They may exhibit, for example, an increased concentration closer to the synapse. The need to reproduce a non-uniform distribution of protein concentration may explain why SAT is bidirectional (in addition to an anterograde component, it also contains a retrograde component).

## 1. Introduction

Because of the great lengths of axons, various axonal cargos are transported in axons utilizing a transport system that is similar to a railway system. Microtubules (MTs) play the role of railway tracks while molecular motors play the role of locomotives pulling axonal cargos along the MT tracks. Kinesin motors move cargos in the anterograde direction (toward the synapse) while dynein motors move cargos in the retrograde direction (toward the soma) [1].

Depending on the average velocity of cargos, axonal transport is divided into fast (characterized by the average cargo velocity ~1 μm s^-1^) and slow axonal transport (SAT). SAT is further divided into slow component-a (SCa), which moves cytoskeletal elements, such as neurofilaments (NFs), with an average anterograde velocity of 0.002–0.02 μm s^-1^ and slow component-b (SCb), which transports ~200 proteins from the soma to the presynaptic terminal with an average anterograde velocity of 0.02–0.09 μm s^-1^ [2–8]. The mechanism of SAT is quite complicated. Diffusion in the cytosol plays an important role in transporting SCb proteins in short axons [9]. On the other hand, in longer axons, molecular motor-driven transport contributes to protein transport in axons [10–12]. The motor-driven components of SAT are propelled by the same molecular motors that move cargos by fast axonal transport: kinesins (anterograde motors) and dyneins (retrograde motors). The difference is that, in the case of SAT, cargos are not moving along MTs continuously, but rather move intermittently, switching between short runs and long pauses, although with anterograde bias [13,14].

The model developed in [12,15] suggests that SAT of SCb proteins is driven by three components: the anterograde kinesin-driven component, the retrograde dynein driven component, and the diffusion-driven component, which describes transport of free (cytosolic) protein. In this paper, we investigate the contribution of these three components of SAT in different segments of the axon: the proximal portion of the axon, the central portion of the axon, and the distal portion of the axon. We use SAT of MAP1B protein as a model system. MAP1B protein is chosen because the distribution of this protein’s concentration along the axon length is reported in [16], the average velocity of SCb transport of MAP1B in the axon is reported in [17], and the diffusivity of MAP1B in the cytosolic state can be easily estimated based on data reported in [9] (see footnote “a” after Table S2). Also, unlike slow axonal transport of tau protein, which includes two diffusion processes (in the cytosol and along microtubules [18,19]), the model of slow axonal transport of MAP1B includes only one mode of diffusion-driven transport (in the cytosol). This makes it easier to analyze the effects of diffusion.

## 2. Materials and models

### 2.1. Mathematical model of MAP1B protein transport in an axon

A neuron with an axon of length *L* was simulated. The coordinate *x* initiates at the axon hillock and is directed toward the axon synapse (Fig. 1a). We simulated MAP1B as an SCb protein [17,20] and assumed that it can exist in five different kinetic states (Fig. 2), as we previously suggested in [21]. An MAP1B concentration in a corresponding kinetic state is given below in parentheses. Two of the kinetic states represent MAP1B undergoing active anterograde (*n_a_*) and retrograde (*n_r_*) transport, which is propelled by kinesin and dynein motors, respectively. Two other kinetic states describe MAP1B in anterograde (*n*_*a*0_) and retrograde (*n*_*r*0_) pausing states, respectively, when motors driving the cargo temporarily disengage from MTs, but are ready to re-engage and resume their motion. In addition to the MAP1B protein associated with MTs, there is also a soluble cytosolic pool of MAP1B protein [22,23]. MAP1B protein in the cytosolic pool must be able to diffuse [24]. This justifies simulating MAP1B in a free (cytosolic) state (*n_free_*).

**Fig. 1.**
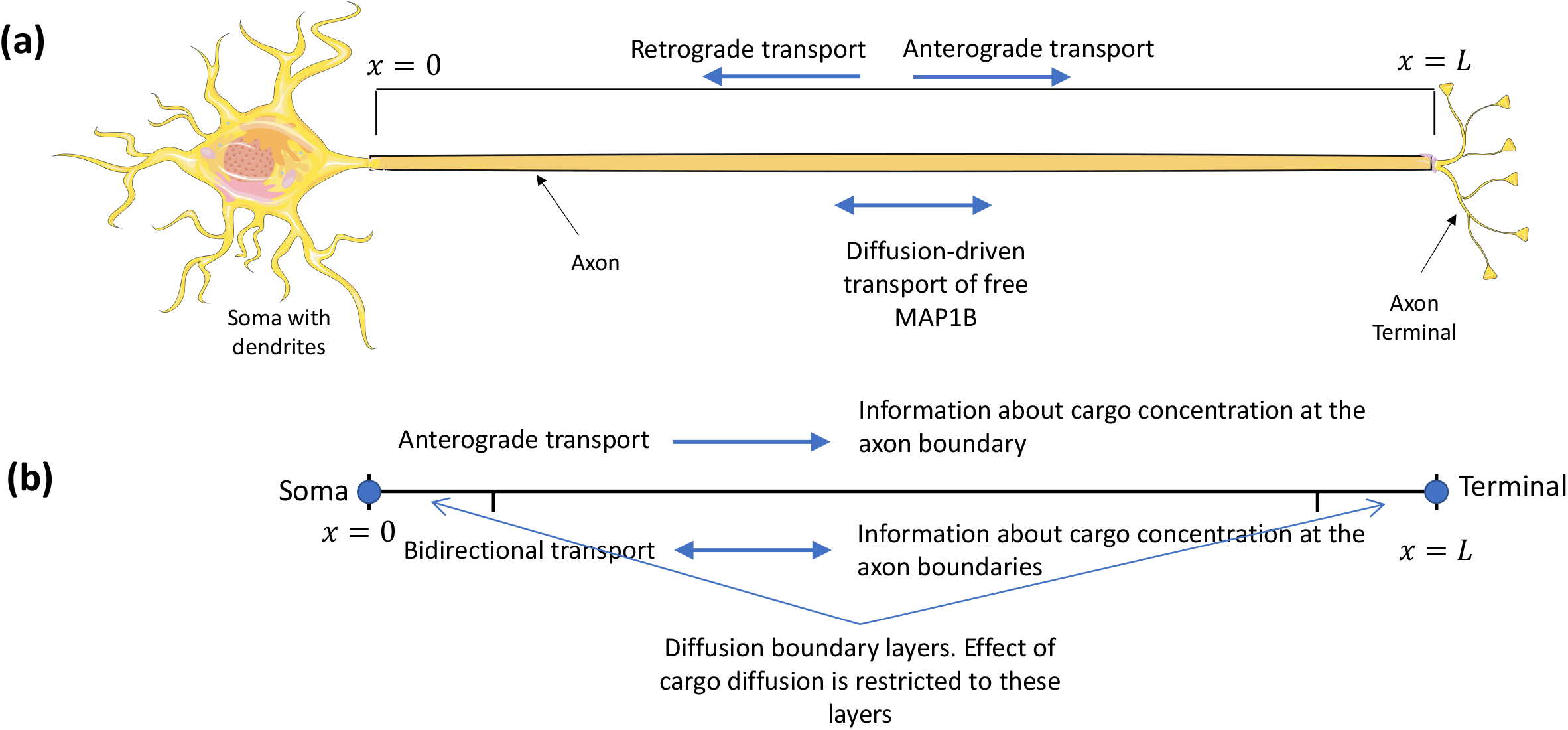
(a) Diagram of a neuron showing various modes of transport of MAP1B protein. Figure is generated with the aid of Servier Medical Art, licensed under a creative commons attribution 3.0 generic license, http://Smart.servier.com. (b) Diagram showing transport of information about the cargo concentration from the axon boundaries in unidirectional and bidirectional axonal transport. To transport cargo against its concentration gradient, the system needs the capability to transport the information about the concentration at *x*=*L* to the soma. By the ability to transport information, we mean how changes in the cargo concentrations at the axon boundaries can be reflected in the distribution of cargo concentration inside the axon.

**Fig. 2.**
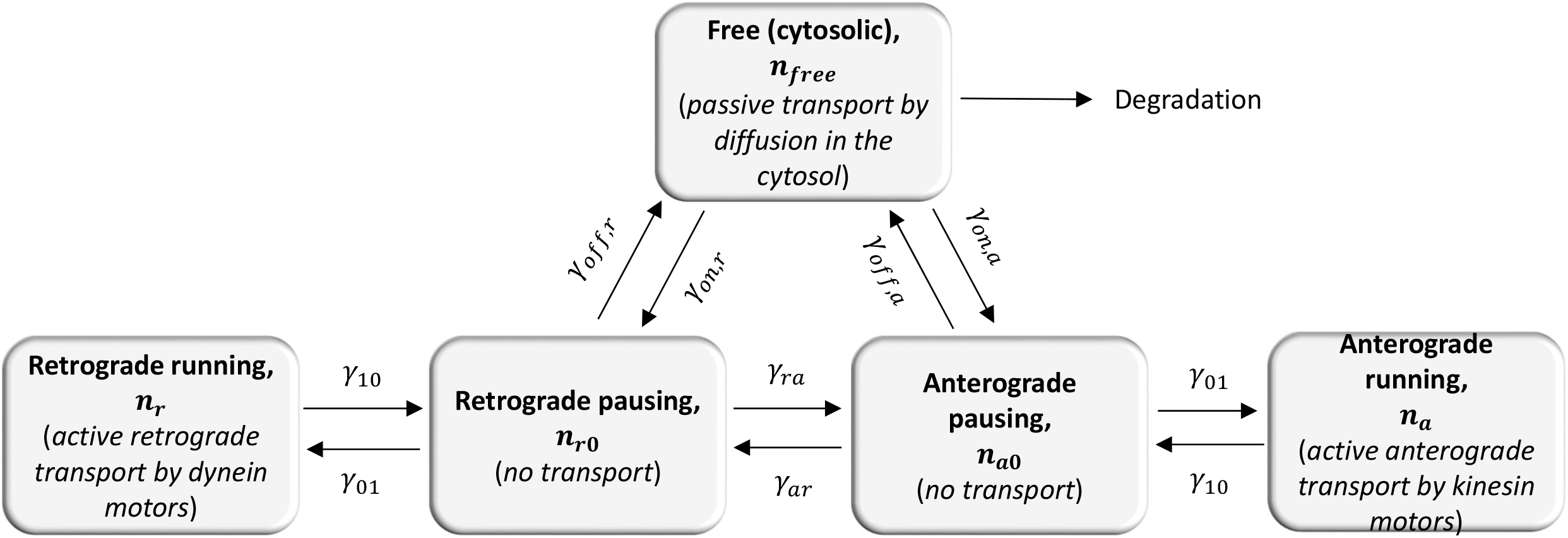
Kinetic diagram showing five kinetic states in our model of MAP1B transport in the axon. Four of these kinetic states are the same as the kinetic states in an SAT model for neurofilaments (see Fig. 4 of [27] and Fig. 1 of [28]).

Model variables are summarized in Table S1, model parameters estimated based on data found in the literature are summarized in Table S2, and model parameters estimated by optimizing the curve fit between model predictions and published experimental data are summarized in Table S3.

The equations expressing MAP1B conservation in the motor-driven states (anterograde and retrograde, respectively) are [21]:

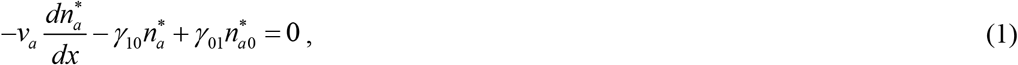

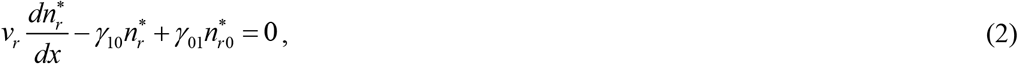

where 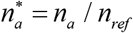 and 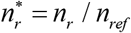. Asterisks denote dimensionless quantities.

Dimensional MAP1B concentrations in various kinetic states are characterized by their linear number density, which is the number of MAP1B protein monomers residing in a particular kinetic state per unit length of the axon (1/μm). Since the governing equations and boundary conditions are linear in terms of MAP1B concentrations, the concentrations can be divided by any appropriately selected reference value (*n_ref_*). This is useful because the experimental data of [16] report fluorescence per unit volume (which is proportional to the total MAP1B concentration) rather than the absolute value of MAP1B concentration. *n_ref_* is selected as the total MAP1B concentration in the leftmost point in Fig. 3D of [16]. For this reason, the dimensionless total MAP1B concentration, 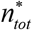, in the leftmost point in Fig. 3a is equal to unity.

**Fig. 3.**
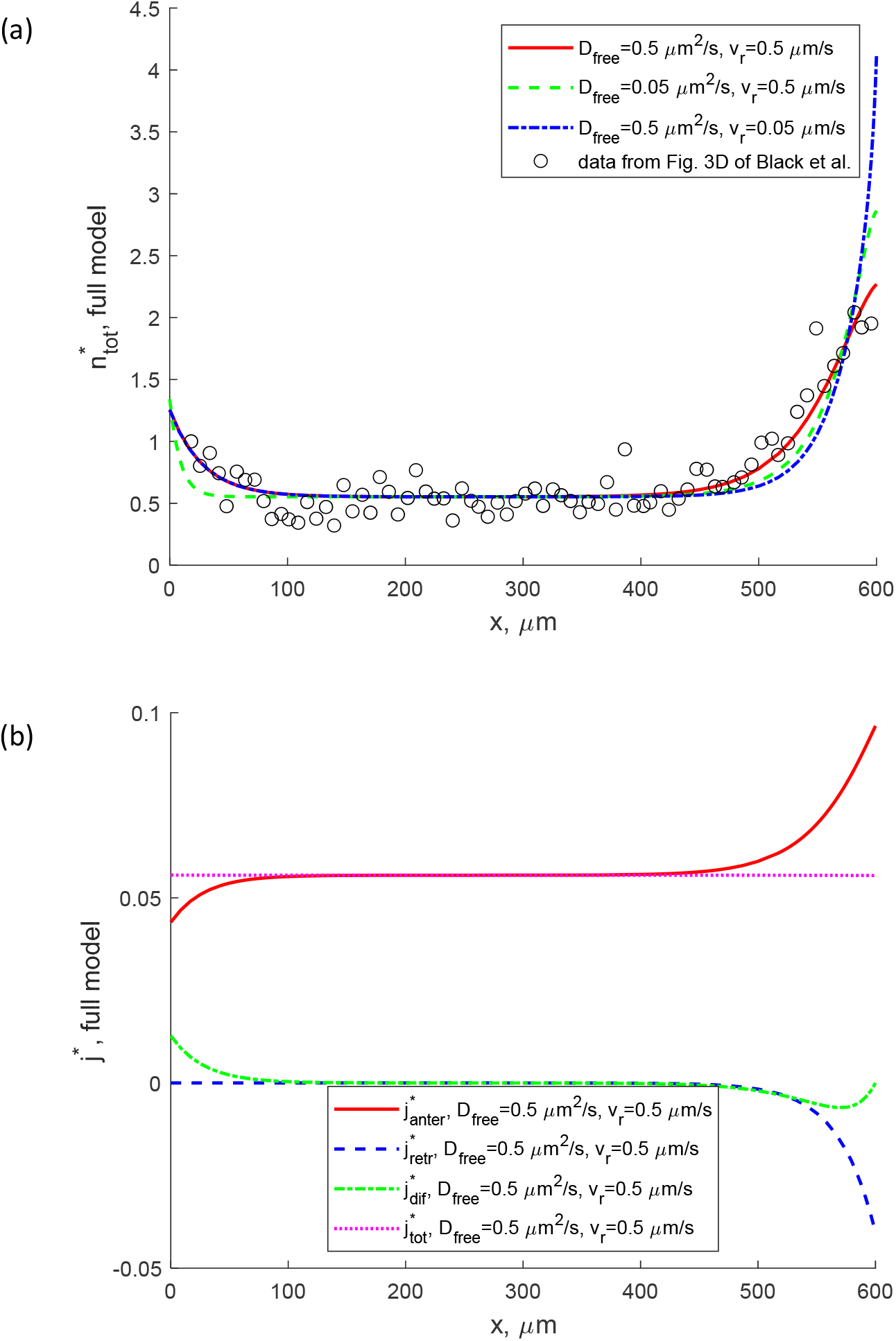
Numerical solution of full SAT model given by Eqs. (1)-(5) with boundary conditions (15a,b) and (16a,b). (a) Total concentration of MAP1B protein; hollow circles show experimental data from Fig. 3D of [16]. Computations are carried out for three different combinations of *D_free_* and *v_r_*. (b) Anterograde (kinesin-driven) flux of MAP1B, retrograde (dynein-driven) flux of MAP1B, diffusion-driven flux of free (cytosolic) MAP1B, and total MAP1B flux (the sum of the three above fluxes). *D_free_* = 0.5 μm^2^/s, *v_r_* = 0.5 μm/s.

The terms in Eqs. (1) and (2) involving the velocities of kinesin and dynein motors, *v_a_* and *v_r_*, respectively, describe the effects of motor-driven transport. The rest of the terms are kinetic terms that describe the effects of MAP1B transition to/from the motor-driven states (see the corresponding arrows in Fig. 2). *γ*_10_ is the kinetic constant describing the probability of MAP1B transition from a running to a pausing state and *γ*_01_ is the kinetic constant describing the probability of MAP1B transition from pausing to a running state (Fig. 2).

The equations stating MAP1B conservation in the pausing states (anterograde and retrograde, respectively) are

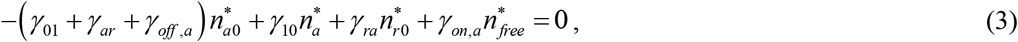

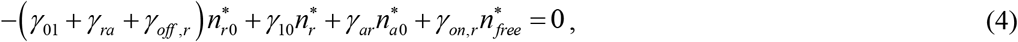

where 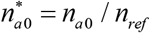 and 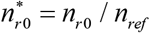. All terms on the left-hand sides of Eqs. (3) and (4) are kinetic terms that describe transitions to/from the pausing states (Fig. 2). *γ_ar_* is a kinetic constant describing the probability of MAP1B transition from the anterograde pausing to the retrograde pausing state, *γ_ra_* is the kinetic constant describing the probability of MAP1B transition from the retrograde pausing to the anterograde pausing state, *γ_on,a_* is the kinetic constant describing the probability of MAP1B transition from the off-track state to the anterograde pausing state, *γ_off, a_* is the kinetic constant describing the probability of MAP1B transition from the anterograde pausing state to the off-track state, *γ_on,r_* is the kinetic constant describing the probability of MAP1B transition from the off-track state to the retrograde pausing state, and *γ_off,r_* is the kinetic constant describing the probability of MAP1B transition from the retrograde pausing state to the off-track state (Fig. 2).

The equation expressing the conservation of free (off-track) MAP1B protein is

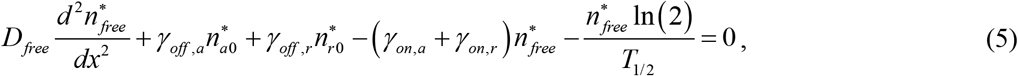

where 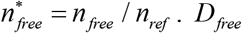. *D_free_* is the diffusivity of MAP1B protein in the off-track state and *T*_1/2_ is the half-life of MAP1B protein. The remaining terms on the left-hand side of Eq. (5) are kinetic terms that describe MAP1B transitions to/from the free cytosolic state (Fig. 2).

The total dimensionless concentration of MAP1B is defined as the sum of concentrations in all five kinetic states (Fig. 2):

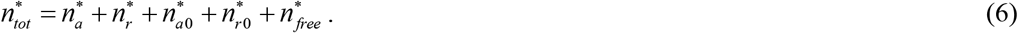

The total flux of MAP1B has contributions from diffusion of free MAP1B as well as from motor-driven transport of this protein in anterograde and retrograde directions, respectively:

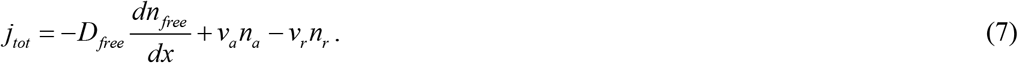

The anterograde kinesin-driven MAP1B flux is defined as:

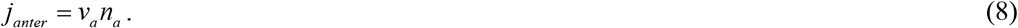

The retrograde dynein-driven MAP1B flux is defined as:

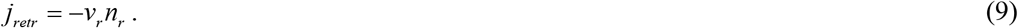

The diffusion-driven flux of free MAP1B is defined as:

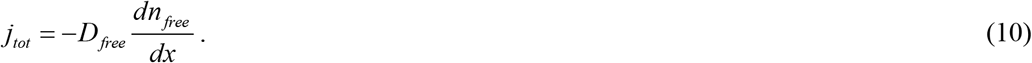

The total dimensionless MAP1B flux is then defined as follows:

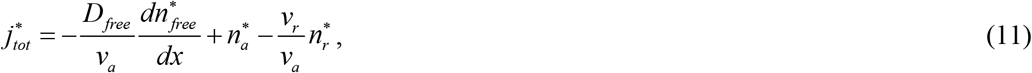

where

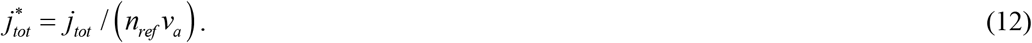

As suggested in [12], the average MAP1B velocity (a quantity that depends on *x*) was then calculated as:

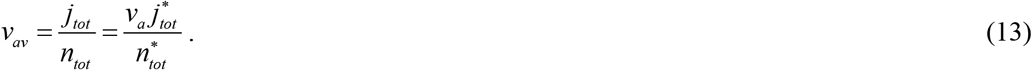

The percentage of MAP1B bound to MTs at a particular location in the axon is found as:

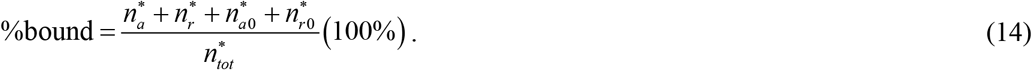

### 2.2. Boundary conditions

Eqs. (1)-(5) were solved subject to the following boundary conditions:

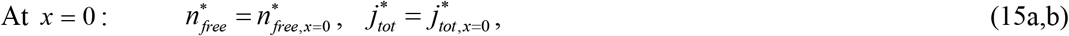

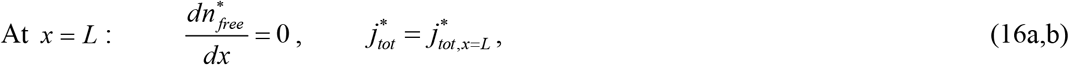

where 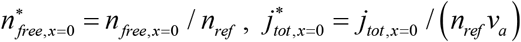, and 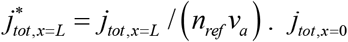 is the total flux of MAP1B into the axon and *n*_*free,x*=0_ is the concentration of free (off-track) MAP1B protein at the axon hillock.

In Eq. (16a), we set the gradient of 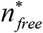 at *x* = *L* equal to zero. There is no physical reason for 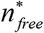 to vary strongly at the end of the axon. Also, computations with other forms of the boundary condition at the axon end caused oscillatory solutions; this behavior contradicts the experimental results of [16]. The derivation of a more detailed form of Eq. (16b) is described in section S1 in Supplementary Materials, see Eq. (S1). The numerical procedure is described in section S2. Values of parameters 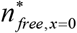 and 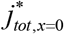, which determine MAP1B concentration and flux at the hillock, respectively, were found by determining the best fit with experimental data (see section S3 and Table S3 in Supplementary Materials).

Note that Eqs. (1)-(5) allow for the imposition of boundary conditions at both ends of the axon, at *x* = 0 and *x* = *L*. This allows for the inclusion of MAP1B concentrations at both ends of the axon in the problem formulation.

### 2.2. Perturbation equations of the MAP1B protein transport model for the cases of small dynein velocity and small diffusivity of free MAP1B protein

#### 2.2.1. Dimensionless retrograde velocity and dimensionless MAP1B protein diffusivity

We are interested in investigating whether the model can simulate cargo transport against the cargo’s concentration gradient, which is a typical situation for many axonal proteins, by kinesin-driven transport alone. This requires setting the dynein velocity and the MAP1B diffusivity to small values. For this purpose, the following small parameters were introduced. The dimensionless retrograde velocity was defined as:

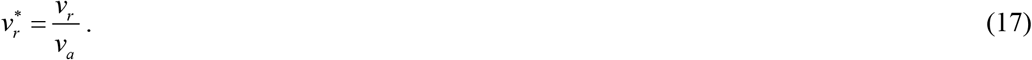

The dimensionless MAP1B diffusivity in the cytosolic state was defined as:

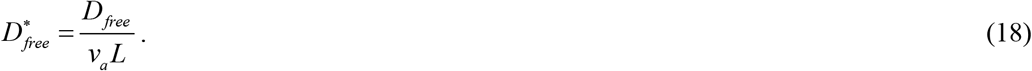

The right-hand side of Eq. (16) represents the reciprocal of the Peclet number. Eq. (16) can be re-written as:

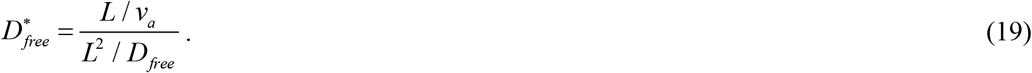

Thus, 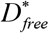 can be viewed as the ratio of the convection time to diffusion time for free MAP1B protein.

#### 2.2.2. Perturbation equations for the case of small dynein velocity, 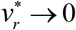

If the dynein velocity is small, 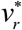 can be treated as a small parameter. For five dependent variables of the model, 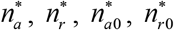, and 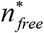, we used the following perturbation expansions (for example):

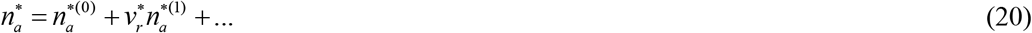

We then substituted the perturbation expansions (such as Eq. (20)) for all five dependent variables into Eqs. (1)-(5). We separated the terms that do and do not contain the small parameter 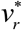 in the obtained equations and equated the terms that do not contain 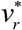 to zero. As a result, Eqs. (S3)-(S7) were obtained.

#### 2.2.3. Perturbation equations for the case of small diffusivity of free MAP1B protein, 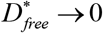

If the diffusivity of cytosolic MAP1B protein is small, 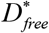 can be treated as a small parameter. For five dependent variables of the model, 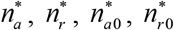, and 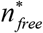, we used the following perturbation expansions (for example):

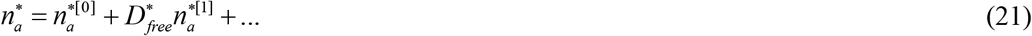

We then substituted the perturbation expansions (such as Eq. (21)) for all five dependent variables into Eqs. (1)-(5). We separated the terms that do and do not contain the small parameter *D_free_* in the obtained equations and equated the terms that do not contain 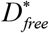 to zero. As a result, Eqs. (S10)-(S14) were obtained.

#### 2.2.4. An analytical solution for the case when both dynein velocity and diffusivity of free MAP1B protein are small, 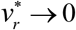 and 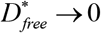

We treated 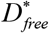 as a small parameter and used the following perturbation expansions for five dependent variables in Eqs. (S3)-(S7), 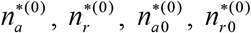, and 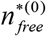. For example, we used the following perturbation expansions for 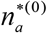:

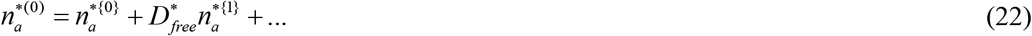

We then substituted the perturbation expansions (such as Eq. (22)) for all five dependent variables into Eqs. (S3)-(S7). We separated the terms that do and do not contain the small parameter *D_free_* in the obtained equations and equated the terms that do not contain 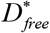 to zero. As a result, Eqs. (S17)-(S21) were obtained.

Because Eqs. (S17)-(S21) allow for the imposition of only one boundary condition (Eq. (S22)), which prescribes the flux of cargo that enters the axon, for 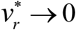 and 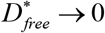 the model does not allow the imposition of a boundary condition describing the cargo concentration at the synapse. This means that for the case of 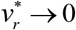 and 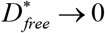 the model does not have the ability to simulate a prescribed distribution of cargo concentration in the axon because it cannot require the concentration of cargo to reach a prescribed value at the synapse. This supports the explanation first proposed in [25], where we suggested that SAT is bidirectional to allow for the maintenance of concentration gradients along the axon length; unidirectional anterograde transport of large cargos that have small diffusivity cannot support concentration gradients (Fig. 1b).

We eliminated 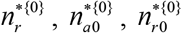, and 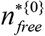 from Eqs. (S17)-(S21), and integrated the obtained equation subject to the boundary condition given by Eq. (S22). This gave the following solution for 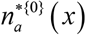:

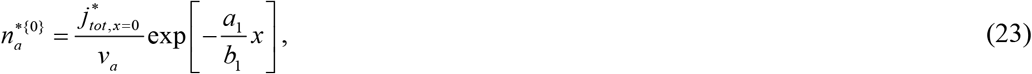

where

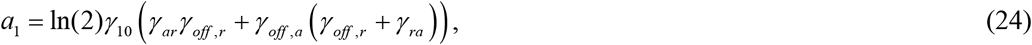

and

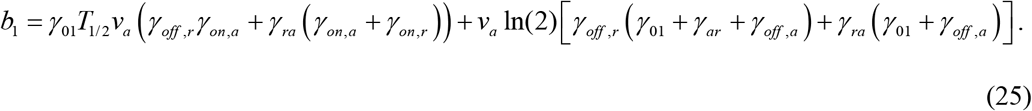

By eliminating, 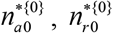, and 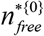 from Eqs. (S18)-(S21), and solving the obtained equation for 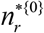, the following was obtained:

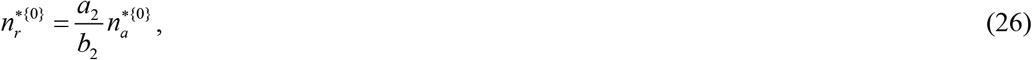

where

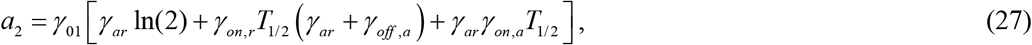

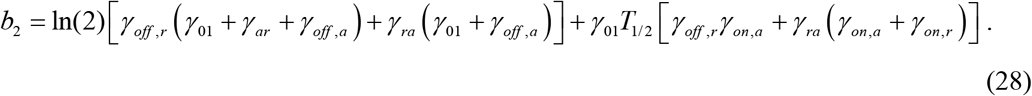

By eliminating, 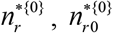, and 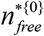 from Eqs. (S18)-(S21), and solving the obtained equation for 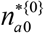, the following was obtained:

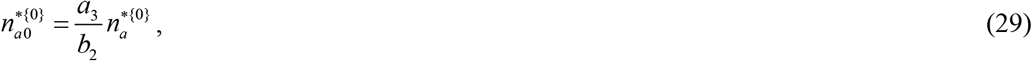

where

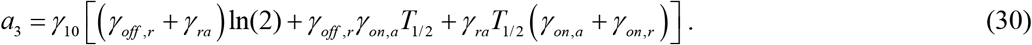

By eliminating, 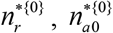, and 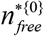 from Eqs. (S18)-(S21), and solving the obtained equation for 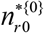, the following was obtained:

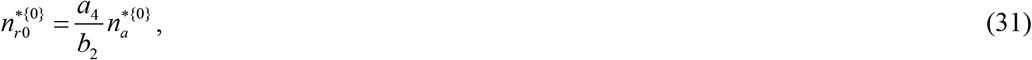

where

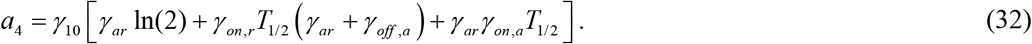

By eliminating, 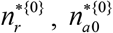, and 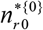 from Eqs. (S18)-(S21), and solving the obtained equation for 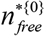, the following was obtained:

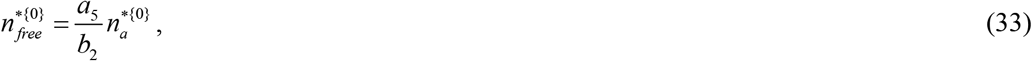

where

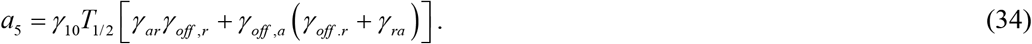

## 3. Results

Model parameters given in Table S3 were determined by minimizing the penalty function given by Eq. (S2). The method is described in section S3 in Supplementary Materials. The first term in the penalty function characterizes the difference between the MAP1B concentration experimentally measured in [16] and the MAP1B concentration numerically predicted by our model. The fitting was done for *v_r_* = 0.5 μm/s and *D_free_* = 0.5 μm^2^/s. Therefore, for these values of the dynein velocity and MAP1B diffusivity the model prediction (solid red line) perfectly fits the experimental data (Fig. 3a). For all other computed cases, we kept the values of parameters given in Table S3 unchanged. This is done to investigate the effect of only one parameter at a time on distributions of the MAP1B concentration and flux.

A decrease of MAP1B diffusivity by a factor of 10 (to *D_free_* = 0.05 μm^2^/s) while maintaining the same value for *v_r_* (0.5 μm/s) results in a reduction of the thickness of two boundary layers, which are located at *x* = 0 and *x* = *L* ends of the axon (Fig. 3a, see also the diagram in Fig. 1b). This suggests that diffusion affects SAT only in the diffusion boundary layers at the axon hillock and axon synapse. It does not affect MAP1B transport over most of the axon except in these boundary layers. This is further confirmed by the distribution of the diffusion-driven flux of MAP1B in Fig. 3b. The diffusion-driven flux is equal to zero over most of the axon, except in the diffusion boundary layer at the axon hillock, where the diffusion-driven flux is positive and thus helps driving MAP1B from the soma into the axon, and the diffusion boundary layer at the axon tip, where the diffusion-driven flux is negative. At the axon tip, *x* = *L*, the diffusion-driven flux is equal to zero (Fig. 3b), which is explained by the boundary condition (16a). The retrograde flux is zero everywhere in the axon except in the boundary layer at the axon tip, which we called the boundary layer of retrograde velocity (Fig. 3b). This is explained by the boundary condition at the axon tip (Eq. (S1)), which assumes that some of MAP1B turn around at the synapse by changing kinesin to dynein motors and travels back toward the soma by retrograde transport. The anterograde flux stays constant over most of the axon except in the boundary layers at the axon hillock and the axon tip. The total MAP1B flux is constant and uniform along the whole axon, which is explained by the conservation of MAP1B at steady-state. There is a very slight decline in the total flux (so small that it is invisible in Fig. 3b), which is caused by a finite half-life of free MAP1B due to its destruction in proteasomes (modeled by the last term on the left-hand side of Eq. (5)).

A reduction of the diffusivity of free MAP1B by a factor of 10 (to *D_free_* = 0.05 μm^2^/s) decreases the thickness of diffusion boundary layers at the axon hillock and the axon tip (Fig. 4a). The thickness of the boundary layer of retrograde velocity is unaffected by the reduction of MAP1B diffusivity (Fig. 4a). A reduction of the dynein velocity by a factor of 10 (to *v_r_* = 0.05 μm/s) decreases the thickness of the boundary layer of retrograde velocity while the thicknesses of the two diffusion boundary layers remain unaffected (Fig. 4b).

**Fig. 4.**
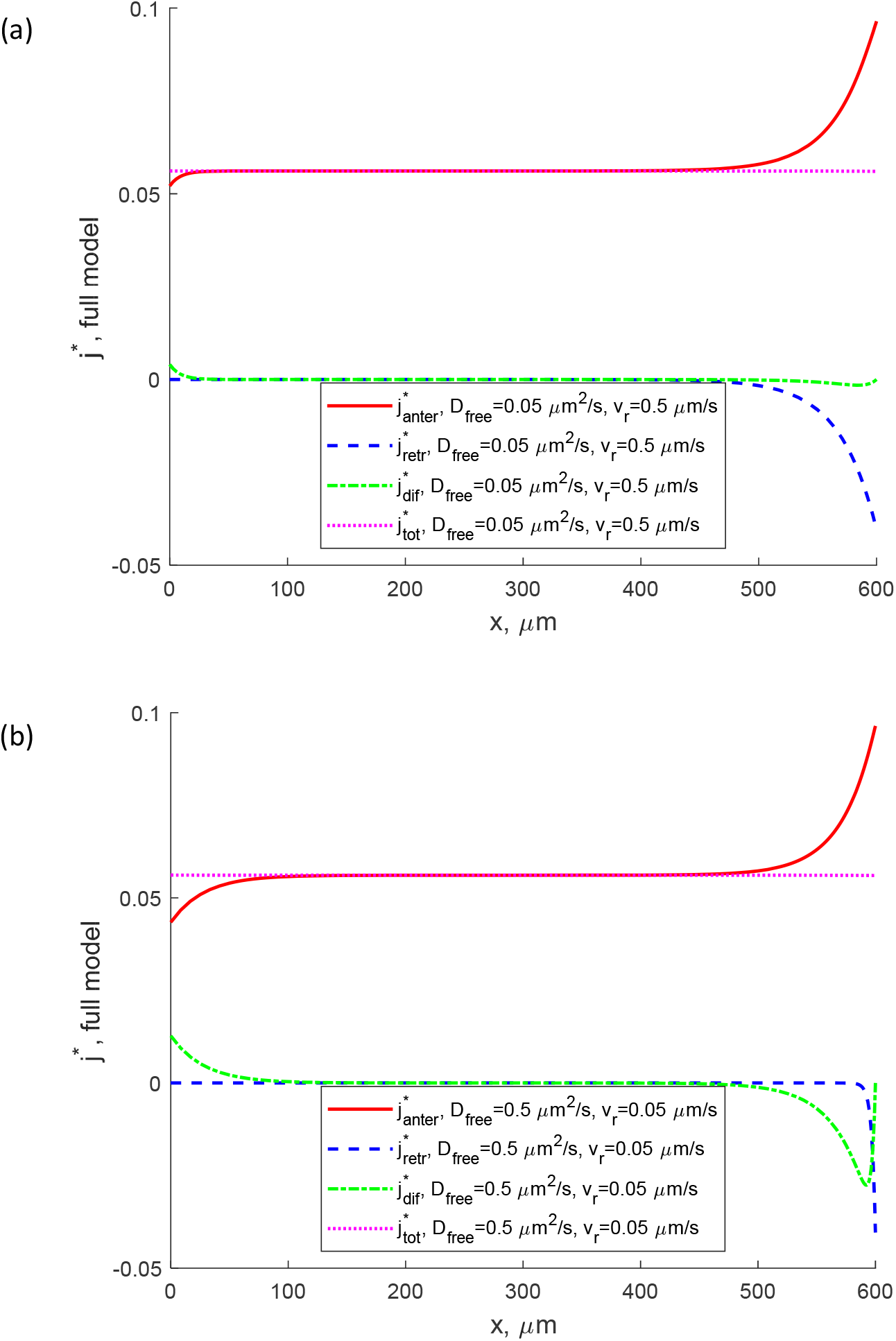
Numerical solution of full SAT model given by Eqs. (1)-(5) with boundary conditions (15a,b) and (16a,b). Anterograde (kinesin-driven) flux of MAP1B, retrograde (dynein-driven) flux of MAP1B, diffusion-driven flux of free (cytosolic) MAP1B, and total MAP1B flux (the sum of the three above fluxes). (a) *D_free_* = 0.05 μm^2^/s, *v_r_* = 0.5 μm/s. (b) *D_free_* = 0.5 μm^2^/s, *v_r_* = 0.05 μm/s.

The observations about the effects of the decrease of *D_free_* and *v_r_* on the thicknesses of diffusion boundary layers and the boundary layer of retrograde velocity are confirmed by investigating distributions of the components of the total MAP1B concentration (Fig. S1). The diffusion boundary layers are characterized by an increased concentration of free MAP1B, 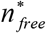. A decrease of *D_free_* decreases the thicknesses of the diffusion boundary layers (Fig. S1b). The boundary layer of retrograde velocity is characterized by the increased concentration of retrograde dynein-driven MAP1B, 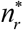. A decrease of *v_r_* decreases the thicknesses of the boundary layer of retrograde velocity (Fig. S1c).

If the dynein velocity is set to zero, the curve fit between model predictions and experimental data becomes less perfect, even for *D_free_* = 0.5 μm^2^/s (Fig. 5a). A decrease of MAP1B diffusivity results in thinner diffusion boundary layers at *x* = 0 and *x* = *L* (Fig. 5b).

**Fig. 5.**
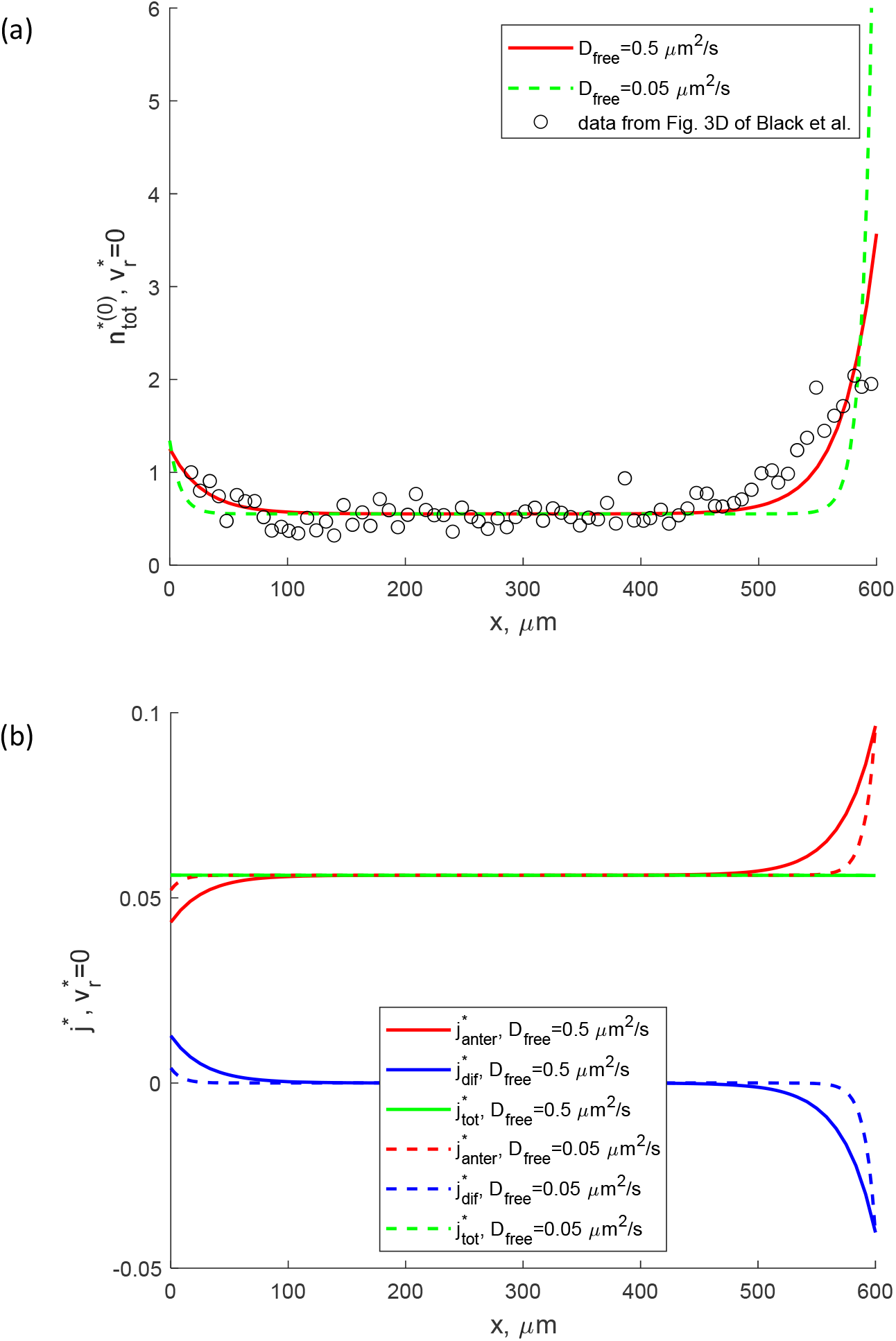
Numerical solution of the perturbation equations (S3)-(S7) for 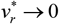 with boundary conditions (S8a,b) and (S9). (a) Total concentration of MAP1B protein for *D_free_* = 0.5 μm^2^/s and *D_free_* = 0.05 μm^2^/s. Hollow circles show experimental data from Fig. 3D of [16]. (b) Anterograde (kinesin-driven) flux of MAP1B, diffusion-driven flux of free (cytosolic) MAP1B, and total MAP1B flux (the sum of the two above fluxes) for *D_free_* = 0.5 μm^2^/s and *D_free_* = 0.05 μm^2^/s.

A sharp increase in the concentration of free MAP1B at the axon tip when the diffusivity of free MAP1B is decreased from 0.5 μm^2^/s (Fig. S2a) to 0.05 μm^2^/s (Fig. S2b) is explained by the accumulation of MAP1B which turns around at the terminal. When the diffusivity is reduced, a greater concentration gradient is needed to transport MAP1B, which turns around at the synapse and moves away from the axon tip.

If the diffusivity of free MAP1B is set to zero, the diffusion-driven transport of MAP1B equals zero. This explains the lack of diffusion boundary layers in Figs. 6a and 6b. However, a boundary layer of retrograde velocity at the axon tip becomes thinner when the dynein velocity is decreased (Figs. 6a and 6b).

**Fig. 6.**
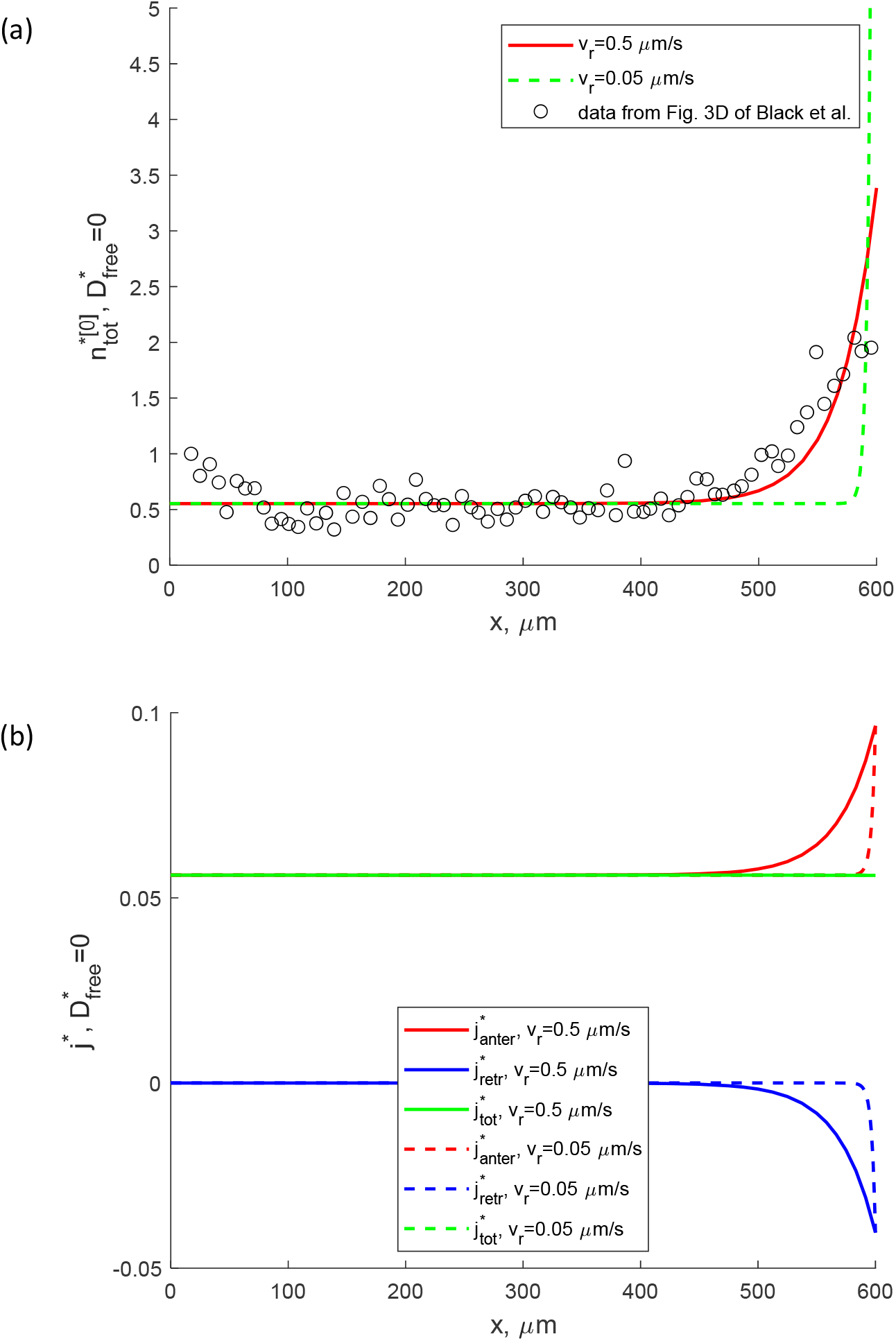
Numerical solution of the perturbation equations (S10)-(S14) for 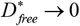 with boundary conditions (S15) and (S16). (a) Total concentration of MAP1B protein for *v_r_* = 0.5 μm/s and *v_r_* = 0.05 μm/s. Hollow circles show experimental data from Fig. 3D of [16]. (b) Anterograde (kinesin-driven) flux of MAP1B, retrograde (dynein-driven) flux of MAP1B, and total MAP1B flux (the sum of the two above fluxes) for *v_r_* = 0.5 μm/s and *v_r_* = 0.05 μm/s.

It should be noted that 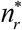 is not the largest concentration in the axon (Fig. S3). The concentrations of MAP1B in different kinetic states at steady-state are determined by equilibrium relations caused by MAP1B transitions between different kinetic states (Fig. 2). The increase of the MAP1B concentration at the axon tip becomes larger when *v_r_* is decreased to 0.05 μm/s (compare Fig. S3a and S3b). This occurs to sustain retrograde transport of MAP1B that turned around at the synapse.

If kinesin-driven anterograde transport is the only component of transport retained in the model (both *v_r_* and *D_free_* are set to zero), the model predicts almost uniform distributions of the total MAP1B concentration (Fig. 7a). All five components of the total MAP1B concentration also exhibit almost uniform distributions (Fig. 8). The anterograde kinesin-driven flux (which in this case equals to the total MAP1B flux) is also almost uniform (Fig. 7b). A very slight decrease of concentrations and the flux (invisible in Figs. 7 and 8) is due to finite MAP1B half-life in the free (cytosolic) state caused by MAP1B destruction in proteasomes (the last term on the left-hand side of Eq. (5)).

**Fig. 7.**
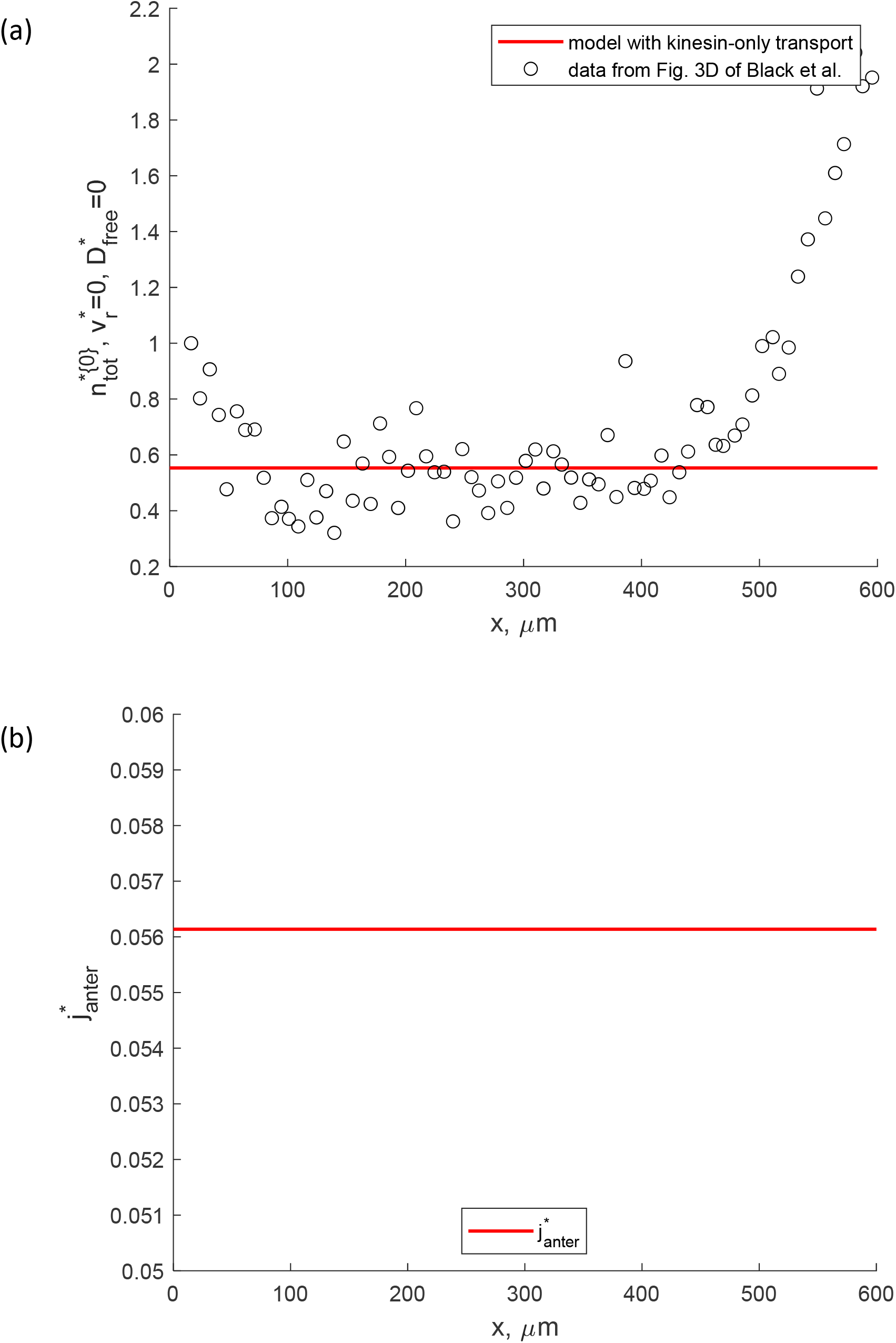
Analytical solution of the perturbation equations (S17)-(S21) for 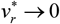 and 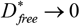 with boundary condition (S22). (a) Total concentration of MAP1B protein; hollow circles show experimental data from Fig. 3D of [16]. (b) Anterograde (kinesin-driven) flux of MAP1B.

**Fig. 8.**
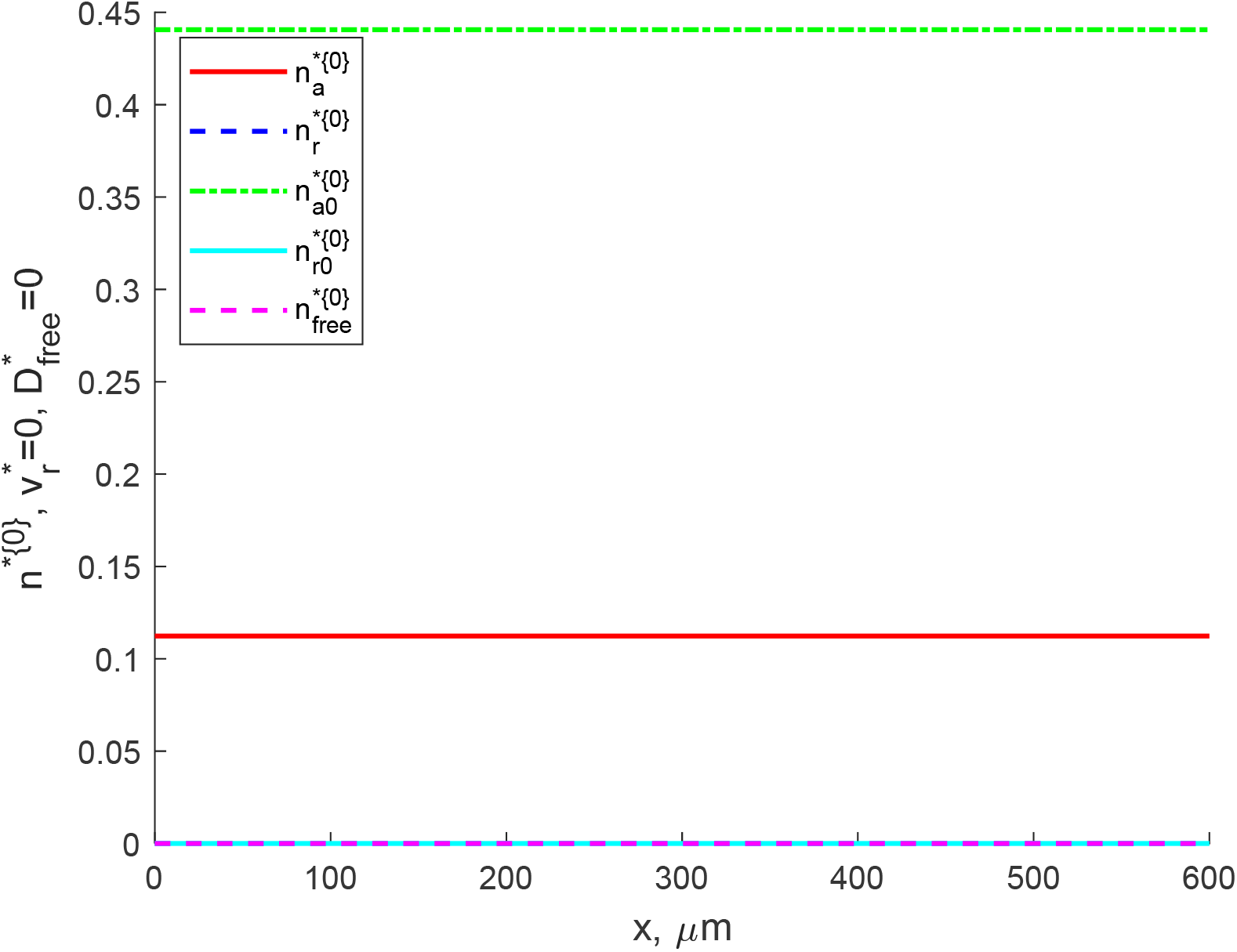
Analytical solution of the perturbation equations (S17)-(S21) for 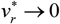 and 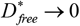 with boundary condition (S22). (a) Concentrations of anterograde kinesin-driven 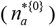, retrograde dynein-driven 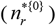, anterograde pausing 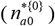, retrograde pausing 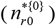, and free 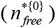 MAP1B protein.

## 4. Discussion and limitations of the model

By analyzing distributions of MAP1B concentrations, we established that SAT of proteins in an axon involves two types of boundary layers. The first type is diffusion boundary layers, which appear at the ends of the axon (at the axon hillock and axon synapse). The two-diffusion-boundary-layer-structure in the tau protein concentration distribution in the axon was first reported in [26]. The thickness of the diffusion boundary layers decreases with a decrease in diffusivity of free (cytosolic) MAP1B but is unaffected by a change in dynein velocity. Since diffusion-driven MAP1B flux is not-zero only within the diffusion boundary layers, we conclude that diffusion contributes to MAP1B transport only near the axon hillock and synapse but does not contribute to MAP1B transport over most of the axon between the two diffusion boundary layers.

The second type is the boundary layer of retrograde velocity, which is located at the axon tip. The retrograde flux is zero everywhere in the axon except in this boundary layer. A decrease in the dynein velocity decreases the thickness of the boundary layer of retrograde velocity; the thickness of this boundary layer is unaffected by the change of MAP1B diffusivity.

Kinesin-driven anterograde transport alone (in the absence of diffusion-driven and dynein-driven transport) cannot explain a variation of MAP1B concentration along the axon length. If all modes of MAP1B transport except for kinesin-driven transport are removed from the model, the model predicts a uniform distribution of the total MAP1B concentration and all of its components (such as anterogradely transported MAP1B, etc.). Since diffusion-driven transport is effective only for protein transport at short distances, the presence of a retrograde dynein-driven component is required to enable a variation of MAP1B concentration along the axon length. This may explain why SAT is bidirectional (it includes both anterograde and retrograde components). Our results suggest that retrograde (dynein) motor dysfunction may lead to neurodegenerative diseases associated with the failure of cargo transport to axon terminals. This may have broad applications to treatment and management of neurodegenerative conditions.

Our model of MAP1B transport is a simplified one as the dynamics of microtubules, polarity and arrangement of microtubules, isoforms of various kinesin/dynein motors, and interactions of MAP1 with other cytoskeletal components are not taken into account. More comprehensive models should be developed in future research. More dedicated experimental data will be needed for the development of such models.

## Data accessibility

This article has no additional data.

## Authors’ contributions

IAK and AVK contributed equally to performing computational work and article preparation. Both authors approved the final version of the manuscript and agreed to be accountable for all aspects of the work.

## Supplementary Materials

### S1. Derivation of the detailed form of Eq. (16b)

A more detailed form of Eq. (16b) was obtained using the same approach that was used in [28] to simulate NF transport. The time it takes for MAP1B protein to turn around at the terminal can be estimated as 1/ *γ_ar_*. Then the probability of MAP1B being degraded at the terminal is 1 – exp [-log (2) / (*γ_ar_T*_1/2_)]. Since the flux of MAP1B that goes into the terminal is *v_a_ n_a_*, the number of MAP1B particles that are degraded in the terminal per second is 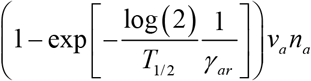. This means that Eq. (16b) can be re-written as:

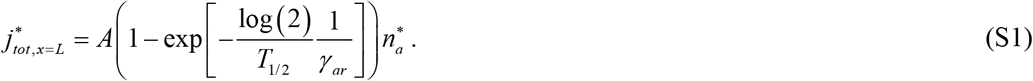

Since we only estimated the rate of MAP1B degradation at the terminal, the left-hand side of Eq. (S1) is multiplied by a dimensionless constant *A* whose value is determined by fitting model predictions with experimental results.

### S2. Numerical solution of differential equations

Since healthy axons operate under the same conditions for many decades, Eqs. (1)-(5) were solved for an axon under steady-state conditions. Eqs. (3) and (4) are algebraic equations, while Eqs. (1), (2), and (5) are ordinary differential equations (ODEs). We eliminated *n*_*a*0_(*x*) and *n*_*r*0_(*x*) from Eqs. (1)-(5) by using Eqs. (3) and (4), as described in [21]. We solved the remaining three ODEs for *n_a_* (*x*), *n_r_* (*x*), and *n_free_*(*x*) by utilizing Matlab’s BVP4C solver (Matlab R2020b, MathWorks, Natick, MA, USA). After the ODEs were solved numerically, *n*_*a*0_ (*x*) and *n*_*r*0_ (*x*) were obtained from Eqs. (3) and (4).

### S3. Method used for finding values of parameters in Table S3 that give the best fit with experimental results for MAP1B protein transport

The optimal values of kinetic constants and parameters characterizing fluxes of MAP1B at the boundaries were found by least square regression (LSR) [29]. The best fit parameters were those that minimized a penalty function, which is defined by the following sum of squared residuals:

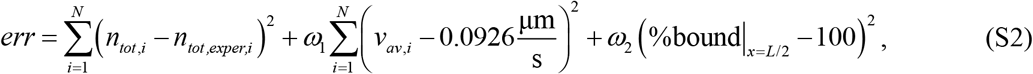

where *ω*_1_ and *ω*_2_ are the weighting factors, which were set to 10 s^2^/μm^2^ and 10, respectively, to provide good visual agreement with experimental data points. We used *N*=76, which is the number of experimental points obtained by digitizing the data reported in [16]. The first term on the right-hand side of Eq. (S2) ensures that the distribution of the total MAP1B concentration predicted by the model is close to the one reported in [16]. The second term constrains the MAP1B transport velocity to approximately 8 mm/day (0.0926 μm/s), which is the average of the range reported in [17]. Ref. [17] reported MAP1B velocity to be between 0.0810 and 0.104 μm s^-1^, which is the upper bound of SCb (ref. [17] called it slow component-c). The third term ensures that most of the MAP1B in the center of the axon is bound to MTs, which may not be the case close to the soma or the terminal because of reactions that MAP1B may be a part of in these locations. The value 10 for the weighting factors *ω*_1_ and *ω*_2_ was found by means of extensive numerical experimentation (data not shown). For example, if *ω*_1_ is set to a very large value, the predicted total MAP1B concentration strays too far from values reported in [16]. A large value of *ω*_1_ would also cause overfit in terms of MAP1B velocity, because it would force the resultant average MAP1B velocity to approach exactly 0.0926 μm/s, ignoring the velocity’s natural variance.

The global minimum was found by using routine MULTISTART with a local solver FMINCON, which can be found in Matlab’s Optimization Toolbox. 5000 randomly selected starting points in the parameter space were used to start the optimization procedure. The best fit values of parameters were then found by iterating from these starting points by using FMINCON [21].

### S4. Zero-order perturbation equations

#### S4.1. Zero-order perturbation equations for 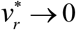 at steady-state

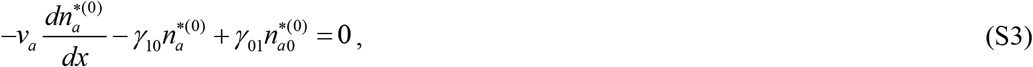

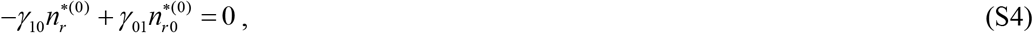

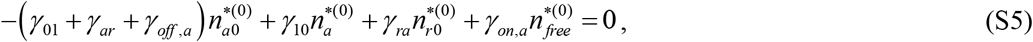

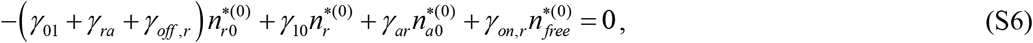

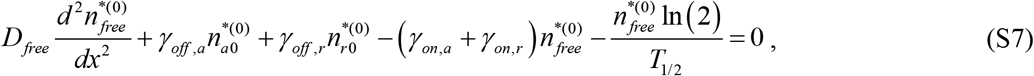

Since the system of Eqs. (S3)-(S7) is of the third order, this system must be solved subject to three boundary conditions. We used the following boundary conditions for Eqs. (S3)-(S7):

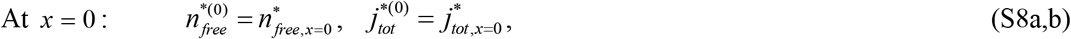

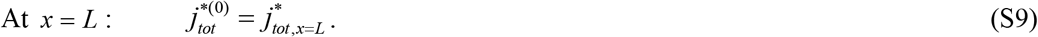

#### S4.2. Zero-order perturbation equations for 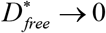 at steady-state

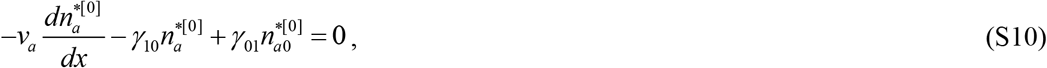

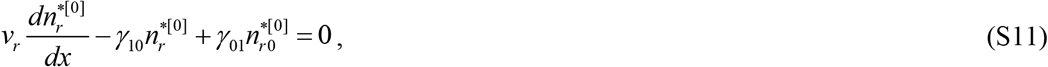

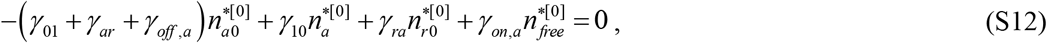

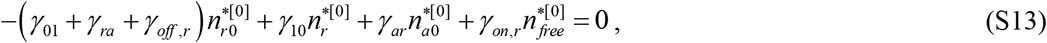

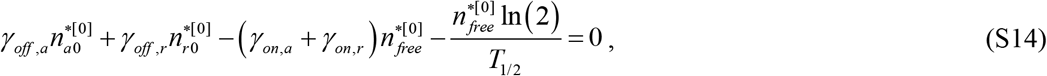

Since the system of Eqs. (S10)-(S14) is of the second order, this system must be solved subject to two boundary conditions. We used the following boundary conditions for Eqs. (S10)-(S14):

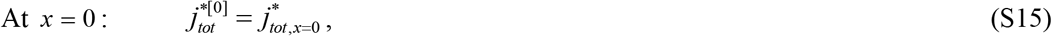

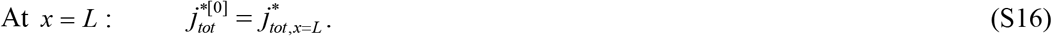

#### S4.3. Zero-order perturbation equations for 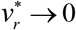 and 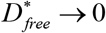 at steady-state

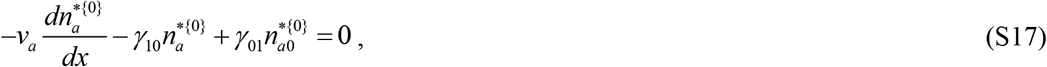

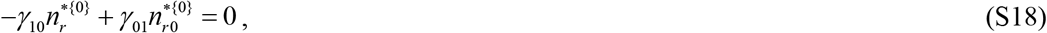

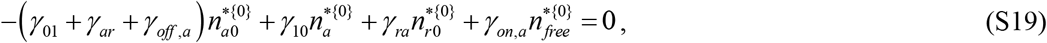

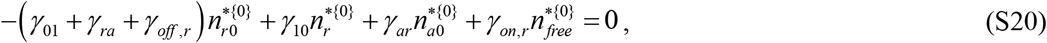

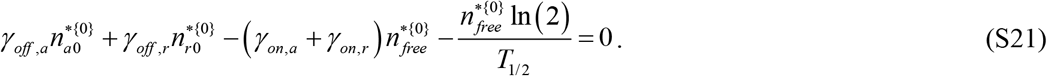

Since the system of Eqs. (S17)-(S21) is of the first order, this system must be solved subject to one boundary condition. We used the following boundary condition for Eqs. (S17)-(S21):

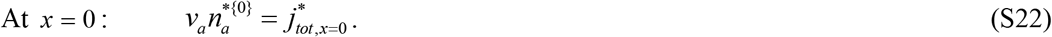

### S5. Supplementary tables

Model variables are summarized in Table S1, model parameters that are found in published literature or estimated based on published results are summarized in Table S2, and model parameters that are found by minimizing the discrepancy between model predictions and published data on MAP1B distribution in the axon and its average velocity are summarized in Table S3.

**Table S1.** Model variables.

| Symbol | Definition | Units |
| --- | --- | --- |
| $n_a^*$ | Dimensionless concentration of on-track MAP1B moving along MTs anterogradely, propelled by molecular motors, $n_a / n_{ref}$ | |
| $n_r^*$ | Dimensionless concentration of on-track MAP1B moving along MTs retrogradely, propelled by molecular motors, $n_r / n_{ref}$ | |
| $n_{a0}^*$ | Dimensionless concentration of pausing on-track MAP1B that is still associated with molecular motors and can resume its anterograde motion, $n_{a0} / n_{ref}$ | |
| $n_{r0}^*$ | Dimensionless concentration of pausing on-track MAP1B that is still associated with molecular motors and can resume its retrograde motion, $n_{r0} / n_{ref}$ | |
| $n_{free}^*$ | Dimensionless concentration of free (off-track) MAP1B diffusing in the cytosol, $n_{free} / n_{ref}$ | |
| $x$ | Cartesian coordinate along the axon | $\mu\text{m}$ |

**Table S2.** Model parameters estimated from the literature.

| Symbol | Definition | Units | Reference | Estimated value |
| --- | --- | --- | --- | --- |
| $D_{free}$ | Diffusivity of MAP1B protein in the off-track state | $\mu\text{m}^2/\text{s}$ | [9]<sup>a</sup> | 0.5 |
| $L$ | Length of the axon | $\mu\text{m}$ | [16] | 600 |
| $T_{1/2}$ | Half-life of MAP1B protein | s | [20] | $5.01 \times 10^5$ |
| $v_a, v_r$ | Velocities of kinesin and dynein motors, respectively | $\mu\text{m/s}$ | [9] | 0.5, 0.5 |

<sup>a</sup> Diffusivity of soluble MAP1B was estimated as follows. The mass of tau protein is between 45-65 kDa, depending on the isoform [30] while the mass of MAP1B is between 320-330 kDa [20]. The diffusivity of the tau protein is approximately $3 \mu\text{m}^2/\text{s}$ [9]. Since according to the Stokes-Einstein equation, the diffusivity of a particle is inversely proportional to its size [31], the diffusivity of soluble MAP1B protein is estimated to be approximately $0.5 \mu\text{m}^2/\text{s}$ .

**Table S3.** Model parameters whose values were determined by finding the values that give the best fit between model predictions and published experimental data (minimizing the objective function *err* defined in Eq. (S2)).

| Symbol | Definition | Units | Estimation method | Estimated value |
| --- | --- | --- | --- | --- |
| $\gamma_{10}$ | Kinetic constant describing the probability of MAP1B transition from the running (anterograde or retrograde) to the corresponding pausing state | $\text{s}^{-1}$ | LSR <sup>a</sup> | $1.600 \times 10^{-2}$ |
| $\gamma_{01}$ | Kinetic constant describing the probability of MAP1B transition from the pausing (anterograde or retrograde) to the corresponding running state | $\text{s}^{-1}$ | LSR <sup>a</sup> | $4.076 \times 10^{-3}$ |
| $\gamma_{ar}$ | Kinetic constant describing the probability of MAP1B transition from the anterograde pausing to the retrograde pausing state | $\text{s}^{-1}$ | LSR <sup>a</sup> | $6.257 \times 10^{-16}$ |
| $\gamma_{ra}$ | Kinetic constant describing the probability of MAP1B transition from the retrograde pausing to the anterograde pausing state | $s^{-1}$ | LSR<sup>a</sup> | $1.453 \times 10^{-8}$ c |
| $\gamma_{on,a}$ | Kinetic constant describing the probability of MAP1B transition from the off-track state to the anterograde pausing state | $s^{-1}$ | LSR<sup>a</sup> | $6.326 \times 10^{-4}$ |
| $\gamma_{on,r}$ | Kinetic constant describing the probability of MAP1B transition from the off-track state to the retrograde pausing state | $s^{-1}$ | LSR<sup>a</sup> | $4.977 \times 10^{-10}$ |
| $\gamma_{off,a}$ | Kinetic constant describing the probability of MAP1B transition from the anterograde pausing state to the off-track state | $s^{-1}$ | LSR<sup>a</sup> | $6.257 \times 10^{-16}$ |
| $\gamma_{off,r}$ | Kinetic constant describing the probability of MAP1B transition from the retrograde pausing state to the off-track state | $s^{-1}$ | LSR<sup>a</sup> | $2.000 \times 10^1$ |
| $j_{tot,x=0}^*$ | Dimensionless total flux of MAP1B into the axon | | LSR<sup>a</sup> | $1.123 \times 10^{-1}$ |
| $n_{free,x=0}^*$ | Dimensionless concentration of free (off-track) MAP1B protein at the axon hillock | | LSR<sup>a</sup> | $7.160 \times 10^{-1}$ |
| $A$ | Coefficient in Eq. (S1) | | LSR<sup>a</sup> | $5.811 \times 10^{-1}$ |

<sup>a</sup> Least square regression, finding parameter values minimizing the penalty function defined by Eq. (S2) by constrained minimization.

### S6. Supplementary figures

**Fig. S1.**
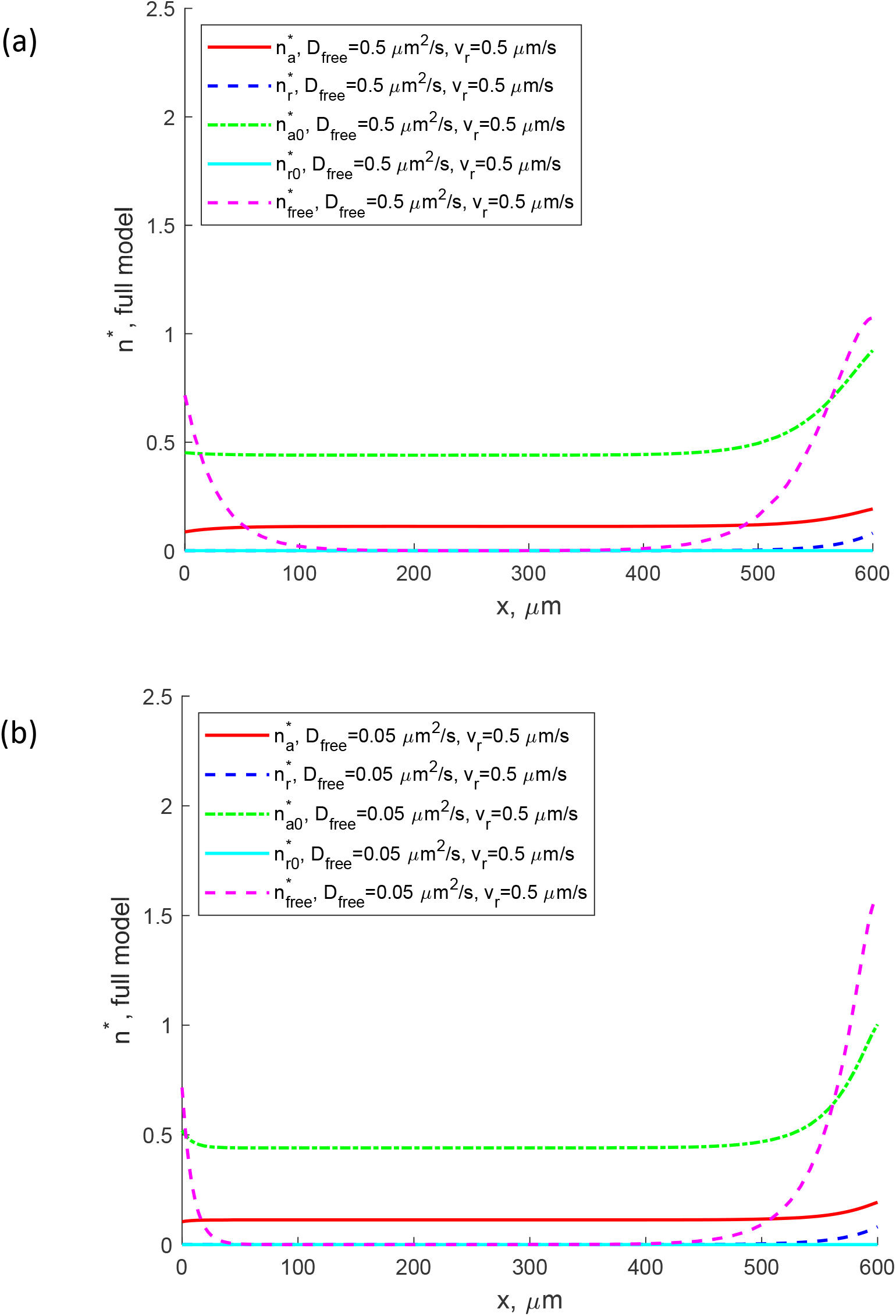
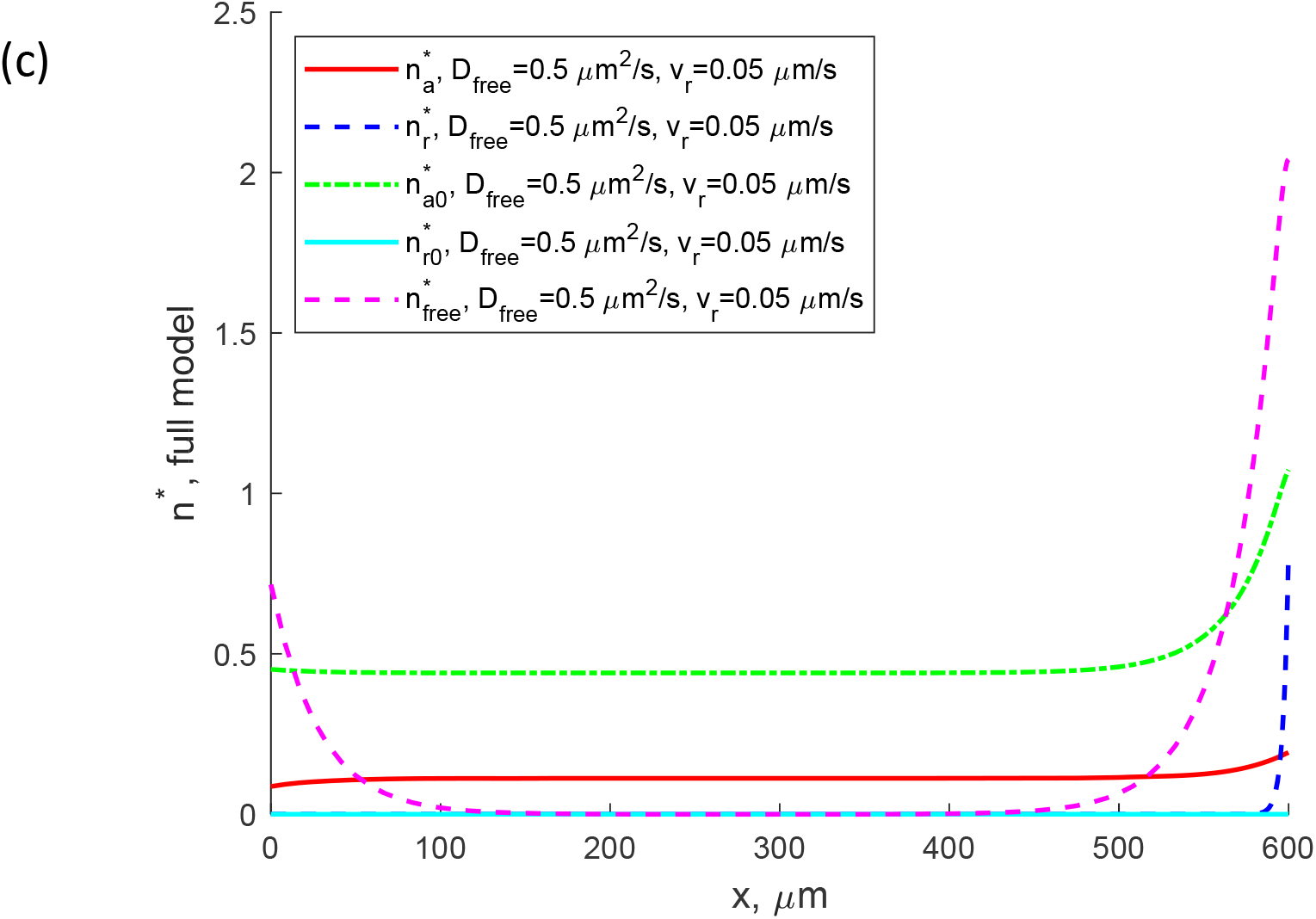
Numerical solution of full SAT model given by Eqs. (1)-(5) with boundary conditions (15a,b) and (16a,b). Dimensionless concentrations of anterograde kinesin-driven 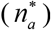, retrograde dynein-driven 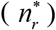, anterograde pausing 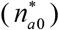, retrograde pausing 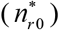, and free 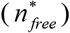 MAP1B protein predicted by the full SAT model. (a) *D_free_* = 0.05 μm^2^/s, *v_r_* = 0.5 μm/s; (b) *D_free_* = 0.5 μm^2^/s, *v_r_* = 0.05 μm/s.

**Fig. S2.**
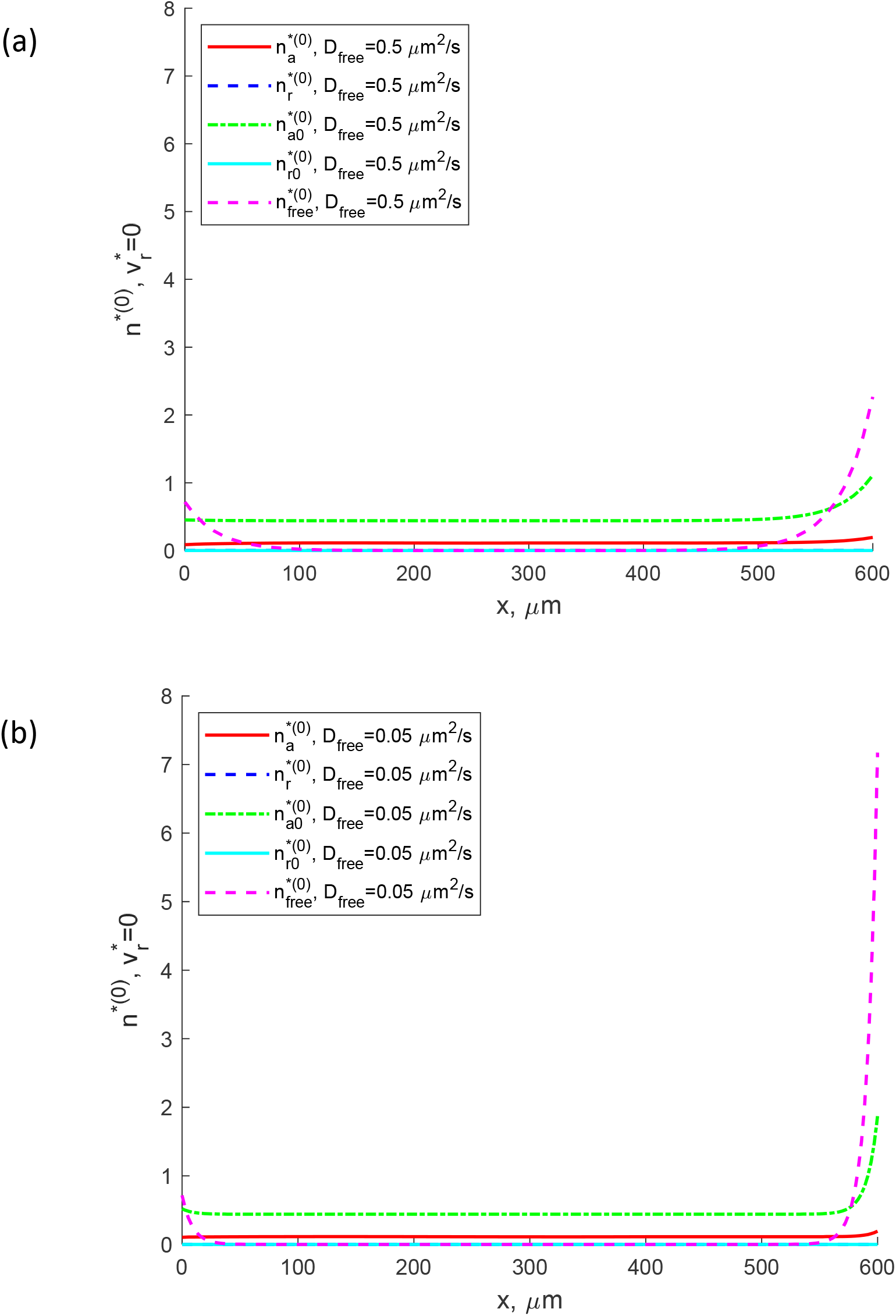
Numerical solution of the perturbation equations (S3)-(S7) for 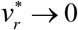 with boundary conditions (S8a,b) and (S9). Concentrations of anterograde kinesin-driven 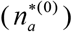, retrograde dynein-driven 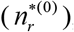, anterograde pausing 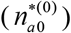, retrograde pausing 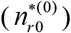, and free 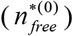 MAP1B protein. (a) *D_free_* = 0.5 μm^2^/s and (b) *D_feee_* = 0.05 μm^2^/s.

**Fig. S3.**
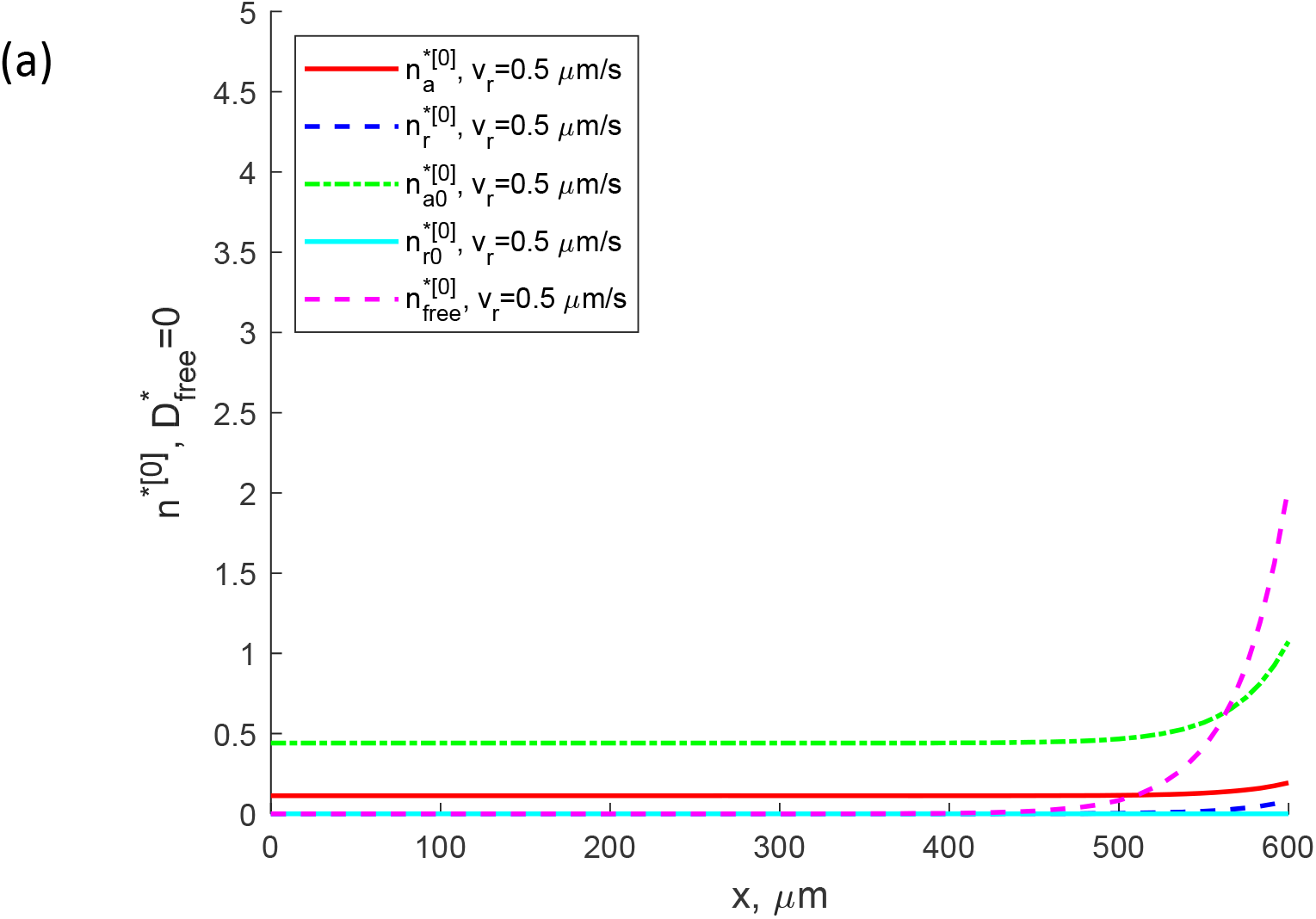
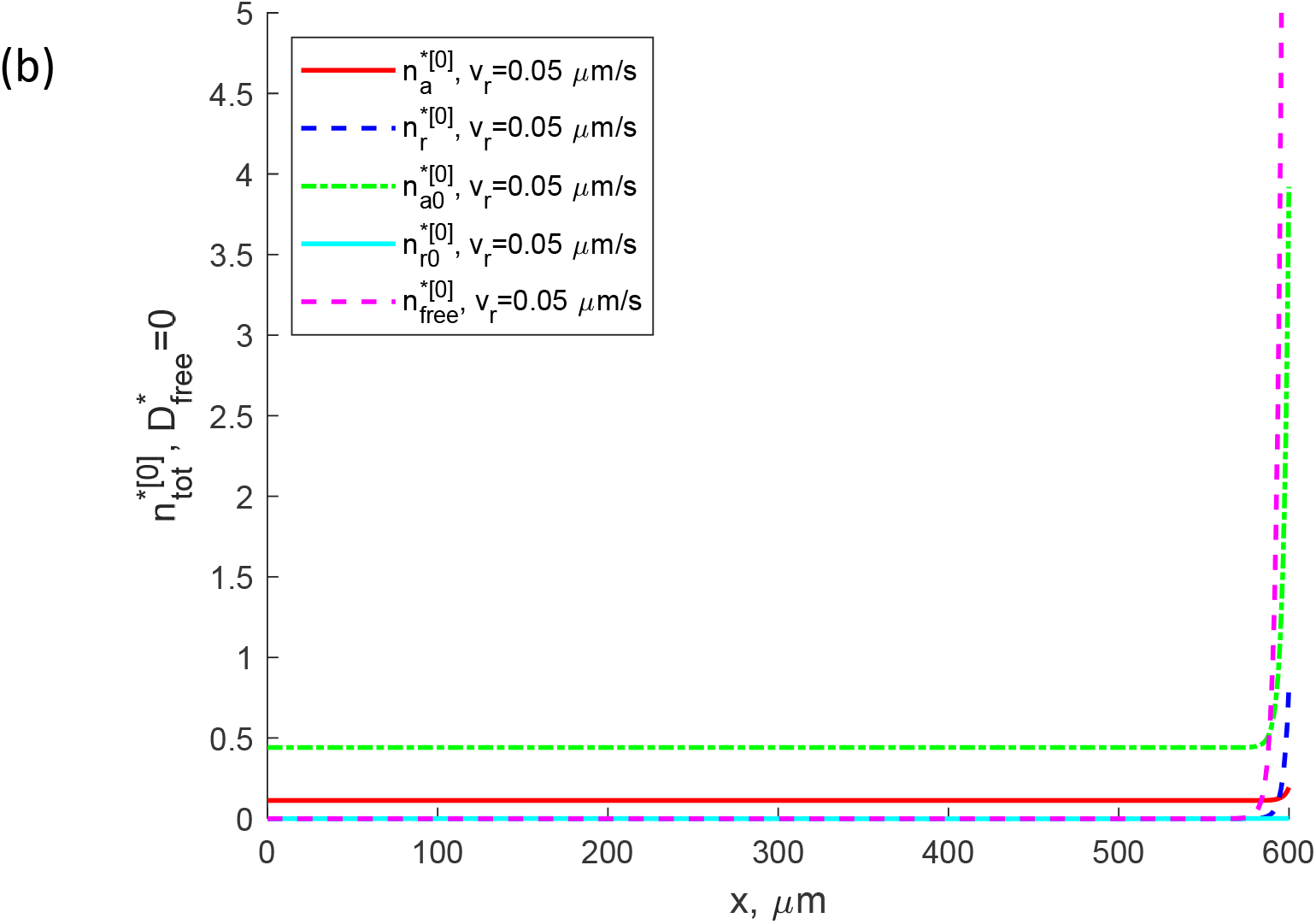
Numerical solution of the perturbation equations (S10)-(S14) for 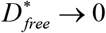 with boundary conditions (S15) and (S16). Concentrations of anterograde kinesin-driven 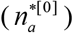, retrograde dynein-driven 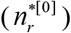, anterograde pausing 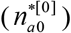, retrograde pausing 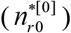, and free 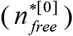 MAP1B protein. (a) *v_r_* = 0.5 μm/s and (b) *v_r_* = 0.05 μm/s.

